# Characterization of the role of glycine lipids in *Bacteroides thetaiotaomicron*

**DOI:** 10.1101/371807

**Authors:** Alli Lynch, Seshu R. Tammireddy, Mary K. Doherty, Phillip D. Whitfield, David J. Clarke

## Abstract

Acylated amino acids function as important components of the cellular membrane in some bacteria. Biosynthesis is initiated by the N-acylation of the amino acid and this is followed by subsequent O-acylation of the acylated molecule resulting in the production of the mature diacylated amino acid lipid. In this study we use both genetics and liquid chromatography-mass spectrometry (LC-MS) to characterize the biosynthesis and function of novel diacylated glycine lipid (GL) species in *Bacteroides thetaiotaomicron*. We, and others, have previously reported the identification of a gene, named *glsB* in this study, that encodes a N-acyltransferase activity responsible for the production of a monoacylated glycine called N-acyl-3-hydroxy-palmitoyl glycine (or commendamide). In all of the *Bacteroidales* genomes so far sequenced the *glsB* gene is located immediately downstream from a gene, named *glsA*, also predicted to encode a protein with acyltransferase activity. We use LC-MS to show that co-expression of *glsB* and *glsA* results in the production of GL in *Escherichia coli*. We constructed a deletion mutant of the *glsB* gene in *B. thetaiotaomicron* and we confirm that *glsB* is required for the production of GL in *B. thetaiotaomicron*. Moreover, we show that *glsB* is important for the ability of *B. thetaiotaomicron* to adapt to stress and colonize the mammalian gut. Therefore, this report is the first to describe the genetic requirements for the biosynthesis of GL, a novel diacylated amino acids species that contributes to fitness in the human gut bacterium, *B. thetaiotaomicron*.

## Introduction

Members of the Phylum Bacteroidetes, including genera containing important human gut commensal bacteria such as *Bacteroides*, *Parabacteroides* and *Prevotella*, dominate the healthy human gut microbiota (1). The gut-associated Bacteroidetes are required to digest complex dietary glycans into short-chain fatty acids (such as acetate and propionate) that are accessible to the host (2-4). A longitudinal study in infants has revealed the presence of *Bacteroides* in the infant gut within 1 week of birth and some species of *Bacteroides* have been shown to utilize the polysaccharides present in human breast milk (5, 6). Therefore it has been suggested that *Bacteroides* may have an important role during the early development of the infant gut (6).

Acylated amino acids can be found in the membranes of many bacteria (7, 8). The best-characterized, and most widespread, acylated amino acid is ornithine lipid (OL). OL contains a 3’-hydroxy fatty acid group attached by an amide linkage to the *α*-amino group of ornithine with a second fatty acid group ester linked to the 3’-hydroxy group of the first fatty acid (9). The genetics of OL production was first described in *Sinorhizobium meliloti* where it was shown that a N-acyltransferase encoded by *olsB* catalysed the attachment of the first fatty acid group to ornithine, resulting in monoacylated ornithine or lyso-OL (10). The second fatty acid was subsequently attached to lyso-OL through the activity of an O-acyltransferase encoded by *olsA*, resulting in the production of OL (11). A bifunctional enzyme called OlsF, with N-terminal homology to OlsA and C-terminal homology to OlsB, has recently been shown to produce OL in *Serratia proteamaculans* (12). Activities that further modify the OL by hydroxylation or methylation have also been identified (13-15). OL have been shown to be important for growth during acid and temperature stress in *Rhizobium tropici* and depletion of OL results in an increase in the speed of crown gall tumor formation in plants infected with *Agrobacterium* (13, 16). Therefore, OL are important during the interactions between bacteria and their environment, including hosts.

Lipid 654 is an acylated serine-glycine dipeptide that has been detected in many members of the Bacteroidetes (17). Lipid 654, also called flavolipin, was first described in members of the *Flavobacterium* and *Cytophaga* (18-20). Some recent studies with *Porphyromonas gingivalis* have implicated Lipid 654 in osteoblast differentiation and atherosclerosis in humans and have also identified Lipid 654 as a potential microbiome-associated biomarker for multiple sclerosis (17, 21-23).

We, and others, have previously identified a gene from *Bacteroides*, initially named *choA*, that encodes a N-acyltransferase required for the production of a monoacylated glycine species, called N-acyl-3-hydroxy-palmitoyl glycine or commendamide (24, 25). In this study we use liquid-chromatography-mass spectrometry (LC/MS) to show that *choA* is also required for the production of a diacylated glycine lipid (GL) in *Bacteroides thetaiotaomicron* and, in line with previous nomenclature, we have renamed *choA* to *glsB*. Using heterologous expression in *E. coli* we show that GL production also requires *glsA*, a gene predicted to encode an O-acyltransferase that is located immediately upstream from *glsB* in the *B. thetaiotaomicron* genome. Finally, we show that *glsB* is important for the ability of *B. thetaiotaomicron* to adapt to stress and colonize the mammalian gut.

## Materials and Methods

### Strains, plasmids, primers and growth conditions

*Bacteroides thetaiotaomicron* VPI 5482 was cultured anaerobically at 37°C in Brain Heart Infusion media (Sigma), supplemented with hemin (5μg ml-1), 0.1% (w/v) cysteine, and 0.2% (w/v) sodium bicarbonate. *Escherichia coli* EPI300 (Epicentre) was routinely cultured in LB broth at 37°C (Merck). For agar plates 1.5% (w/v) agar was added to the liquid media. Where appropriate, antibiotics were added to the media at the following concentrations: Ampicillin (Amp), 100μg/ml; Chloramphenicol (Cm), 12.5μg/ml; Gentamycin (Gm), 50μg/ml or 200μg/ml; Erythromycin (Ery), 25μg/ml. Plasmids and primers used in this study are shown in Supplementary Tables 1 and 2.

### *Construction of gene knock-outs in* B. thetaiotaomicron

Gene deletions were carried out using *B. thetaiotaomicron* Δ*tdk*, as previously described (26). Briefly the DNA flanking the gene to be deleted were amplified and fused by PCR, cloned into the pEXCHANGE-*tdk* vector and transformed into *E. coli* S17-1 *λpir*. The donor (*E. coli*) and recipient (*B. thetaiotaomicron*) strains were mixed, plated onto BHIS agar containing 10% (v/v) horse blood (BHIS blood agar), and incubated, aerobically at 37°C for 24 h. The biomass was re-suspended in 5ml TYG broth, before plating onto BHIS blood agar supplemented with Gm and Ery. The plates were incubated anaerobically for 48 h at 37°C, before 5-10 colonies were re-streaked onto BHIS blood agar (Gm, Ery). After 48 h at 37°C, single colonies were picked into TYG broth and grown for 20 h without antibiotics before plating onto BHIS blood agar supplemented with 200 μg ml^−1^ 5-Fluoro-2’-deoxyuridine (FUdR) for vector counter selection. The plates were incubated anaerobically for 72 h and re-streaked onto BHIS blood + FUdR agar plates. Colony PCR, using primers that were designed outside of the flanking regions, was used to identify potential knock out mutants, before confirmation by sequencing. For complementation experiments the *glsB* (*choA*) gene, plus 500bp upstream from the proposed translation start site, was amplified, and cloned into the pNbu2-bla-ermGb insertion vector. The cloned DNA fragment was inserted into the *B. thetaiotaomicron* Δ*tdk* Δ*glsB* genome into either of the two Nbu2-targetted tRNA^Ser^ loci, via conjugation from *E. coli* S17-1 *λpir*. The resulting complemented strain, *B. thetaiotaomicron* Δ*tdk* Δ*glsB::glsB*, was selected by plating onto BHIS blood agar supplemented with Gm and Ery, and the presence of the *glsB* gene was confirmed by PCR.

### Colonization of germ-free C57BL/6NTac mice

All experiments involving animals were performed at the Biological Services Unit in University College Cork and were approved by the University College Cork Animal Experimentation Ethics Committee. For colonization experiments, 6 week old germ-free female C57BL/6NTac mice were gavaged with 20μl of 5×10^9^ cfu ml^−1^ of the appropriate bacterial strain (n=9 for *B. thetaiotaomicron* WT, n=8 for *B. thetatiotaomicron* Δ*glsB* and n=4 for uninoculated control). The mice were housed as groups of 2-3 in individually-ventilated cages (IVC) and bacterial enumerations were carried out by serial dilution and plating homogenized fecal pellets collected from each IVC on Day 2, 6, 9, and 12 post-gavage. All mice were euthanized on Day 14, the ceca were harvested and cecal contents were collected for further analysis, including bacterial enumeration.

### Analysis of short-chain fatty acids (SCFA)

The level of SCFA in the cecal contents was determined by HPLC using a protocol described previously (27). Cecal contents were weighed and re-suspended in sterile MilliQ water (1:10 (w/v)) containing several 3-4 mm sterile glass beads (Sigma). The samples were vortexed for 1 min and homogenates were centrifuged at 10,000 x *g* for 10 min. The supernatants were filter sterilised using a 0.22 μm filter and analysed using HPLC with a refractive index detector (Agilent 1200 HPLC system). A REZEX 8μ 8%H, Organic Acid Column 300 x 7.8 mM (Phenomenex, USA) was used with 0.01 N H_2_SO_4_ as the elution fluid, at a flow rate of 0.6 ml min^−1^. The temperature of the column was maintained at 65°C and 20 μl of each sample was injected for analysis. End-product peaks were identified by comparison of their retention times with those of pure compounds and concentrations were determined from standards of known concentrations.

### Identification and quantification of glycine lipids

Overnight cultures of *E. coli,* with the appropriate plasmids, or *B. thetaiotaomicron* were inoculated into fresh medium (LB broth with 0.2% (w/v) L-arabinose for *E. coli* or BHIS broth for *B. thetaiotaomicron*) at an OD_600_=0.05 and allowed to grow at 37°C until OD_600_=0.5-0.6. At this point 1 ml samples were centrifuged (5 min, 12,000g) and the pellets were re-suspended in HPLC grade methanol (Sigma) and 500 pmol N-arachidonyl glycine (NAGly 20:4) (Cayman Chemicals, Ann Arbour, MI, USA) was added as an internal standard. Ethyl acetate was added and the mixture was left at 4°C for 30 min before being centrifuged at 2000 x *g* for 5 min to remove denatured proteins. The supernatant was collected, evaporated to dryness under nitrogen gas and reconstituted in methanol containing 5 mM ammonium formate (Sigma). LC-MS analyses were performed using a Thermo Exactive Orbitrap mass spectrometer (Thermo Scientific, Hemel Hempsted, UK) equipped with a heated electrospray ionization (HESI) probe and coupled to a Thermo Accela 1250 ultra-high pressure liquid chromatography (UHPLC) system. Samples were injected on to a Thermo Hypersil Gold C18 column (2.1 mm x 100 mm, 1.9 μm) maintained at 50°C. Mobile phase A consisted of water containing 10 mM ammonium formate and 0.1% (v/v) formic acid. Mobile phase B consisted of 90:10 isopropanol/acetonitrile containing 10 mM ammonium formate and 0.1% (v/v) formic acid. The initial conditions for analysis were 65%A/35%B and the percentage of mobile phase B was increased from 35%-65% over 4 min, followed by 65%-100% over 15 min held for 2 min before re-equilibration to the starting conditions over 6 min. The flow rate was 400 μl/min. Samples were analysed in negative ion mode over the mass to charge ratio (m/z) range 250-2000 at a resolution of 100,000. The signals corresponding to the accurate m/z values for [M-H]-ions of glycine lipid molecular species were extracted from raw LC-MS data sets with the mass error set to 5 ppm. Quantification was achieved by relating the peak area of the glycine lipid species to the peak area of the NAGly 20:4 ISTD. Tandem mass spectrometry (MS/MS) was employed to confirm the identity of glycine lipid species. Samples were infused at a rate of 5 μl/min into a Thermo LTQ-Orbitrap XL mass spectrometer and subjected to higher-energy collision dissociation (HCD) in the Orbitrap analyser. Additional MS^3^ analyses were performed through collision induced dissociation (CID) in the ion trap. Collision energies ranged from 40-65% and helium was used as the collision gas.

### Reverse Transcription PCR (RT-PCR)

Overnight *Bacteroides* cultures were sub-cultured into fresh BHIS broth to an OD_600_=0.05 and incubated anaerobically at 37°C until the cultures reached mid exponential phase (OD_600_=0.3-0.5). At this stage a 5 ml aliquot was removed, centrifuged and the cell pellet re-suspended in RNAprotect (Qiagen). RNA extractions were carried out using High Pure RNA Isolation Kit (Roche), according to manufacturer’s instructions. For the qualitative determination of gene expression, RNA was reverse transcribed into cDNA using QuantiTect Reverse Transcription Kit (Qiagen), according to manufacturer’s instructions. This cDNA was subsequently used as a template for PCR using 0.5 μl DNA template, 2.5 μl CoralLoad PCR buffer (Qiagen), 100 pmol of the appropriate primers (see Supplementary Table 2), 0.2 mM dNTP mix (Promega), 0.125 μl Taq DNA polymerase (Qiagen), and sterile MilliQ dH20, in a final volume of 25 μl. The following PCR conditions were used: 95°C for 5 min (initial denaturation), followed by 35 cycles of 95°C for 30 s, primer specific annealing temperature for 30 s, and 72°C for template specific length of time (1 min per 1kB DNA). This was followed by a final extension of 72°C for 10 min.

### Stress assays

Overnight cultures of the appropriate *B. thetaiotaomicron* strains were adjusted to OD_600_=0.05 in fresh BHIS broth and incubated for 7h at 37°C anaerobically. At this point the OD_600_ was adjusted to OD_600_=0.2 and each culture was split into 3 equal aliquots whereby one aliquot was cultured anaerobically, another aliquot was exposed to air and the final aliquot was incubated in BHIS broth with 1% (w/v) porcine bile (Sigma). Following incubation for 14h under the appropriate stress conditions, viable cells were enumerated by serial dilutions and plating onto BHIS agar.

### Statistical analysis

All statistical analysis was performed using GraphPad Prism 6.0e for Mac software. All experiments were carried out using biological triplicate samples, unless stated otherwise. The Student’s t-test or Mann-Whitney test were used to compare two different groups of data, as indicated. One-way ANOVA, with Tuckey’s post-test, was used to compare three or more groups of data and differences were considered to be significant if *P*<0.05.

## Results

### *ChoA and ChoB are required to produce diacylated GL in* E. coli

We have previously shown that *choA*, when expressed in *E. coli*, results in the production of commendamide, a N-acylated (3-OH C16:0) derivative of glycine with hemolytic activity and the ability to solubilise cholesterol micelles (24). In all members of the Bacteroidales the *choA* gene is located immediately downstream from another gene (nominally called *choB*) predicted to encode an O-acyltransferase (24). Together, *choA* and *choB* are homologous to the bifunctional amino acid acyltransferase, OlsF, that is responsible for the production of OL in *S. proteamaculans* (see Figure 1). Therefore, we wanted to determine whether *choA* and *choB* might work together to produce diacylated glycine lipids (GL). To do this we amplified BVU_RSO7720 *(choA),* BVU_RS07715 *(choB)* and *choB-choA* from *B. vulgatus* and cloned the genes into pBAD24 for arabinose-controlled expression in *E. coli* (resulting in the formation of pBAD-*choA*, pBAD-*choB* and pBAD-*choBA*, respectively). The cells were cultured in LB broth and gene expression was induced by the addition of 0.2% (w/v) L-arabinose to the cultures (as described in Materials and Methods). Cells were harvested and methanol extracts of the cell pellets were subjected to high-resolution LC/MS analysis. In cells overexpressing *choA,* we could detect lipid species that corresponded to a glycine with an N-acyl substitution of varying carbon chain lengths and degrees of saturation, primarily 14:0, 16:0 (i.e. commendamide), 16:1 and 18:1 (see Figure 2 and Table 1). Notably, overexpression of *choA* also resulted in the production of very low levels of diacylated glycine i.e. 0.7% of the total acylated glycine pool (see Table 1). When cells carrying pBAD-*choBA* were analysed we could detect a range of both monoacylated and diacylated glycine species (see Figure 2 and Table 1). The identify of these lipids was confirmed by MS/MS fragmentation although some diacylated glycine species appeared to have mixed fatty acid compositions e.g. in the peak with a m/z value of 510.4 there was a mixture of diacylated glycines substituted with 3-OH-16:0 + 12:0 and 3-OH-14:0 + 14:0 (see Table 1). The total level of monoacylated glycine production in cells overexpressing *choBA* was approximately 4.5-fold lower than the level observed in cells overexpressing *choA* alone. Moreover, diacylated glycine production in *choBA*-expressing cells accounted for 92.7% of the total acylated glycine pool (see Table 1). Importantly, we could not detect monoacylated or diacylated glycine in cells overexpressing *choB* alone (see Figure 2). Therefore, we propose that, in a mechanism analogous to OL biosynthesis, ChoA N-acylates glycine, resulting in the formation of lyso-GL which is, subsequently, O-acylated by ChoB to produce diacylated GL. In accordance with the nomenclature used for the genes involved in OL biosynthesis, we propose to rename *choA* and *choB* as *glsB* (encoding glycine N-acyltransferase) and *glsA* (encoding lyso-GL O-acyltransferase), respectively.

**Figure 1.**
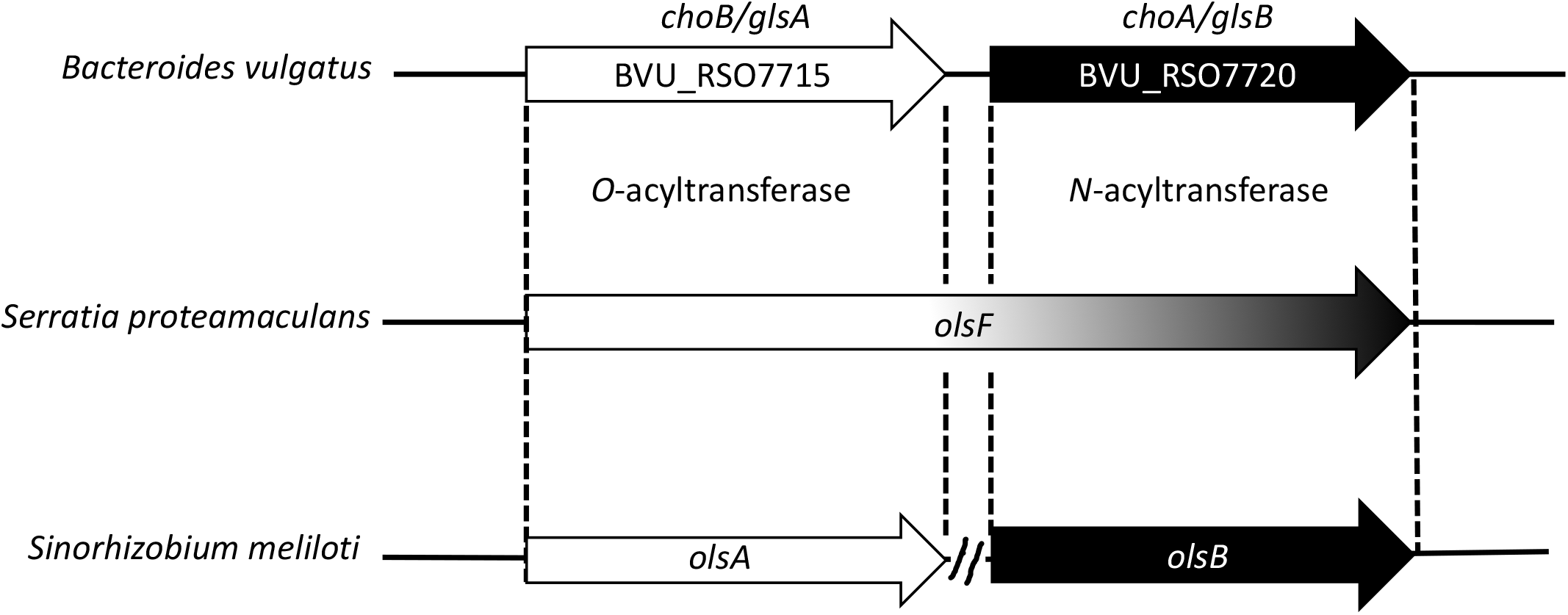
The protein encoded by the *choB* gene in *Bacteroides vulgatus* has predicted homology with OlsA (SM11_chr0038) and the N-terminus of OlsF (Spro_2569), carrying the O-acyltransferase activity required for the biosynthesis of ornithine lipids (OL). Similarly, the *choA* gene is predicted to encode a protein with homology to OlsB (SM11_chr0026) and the C-terminus of OlsF, carrying the N-acyltransferase activity involved in OL biosynthesis.

**Figure 2.**
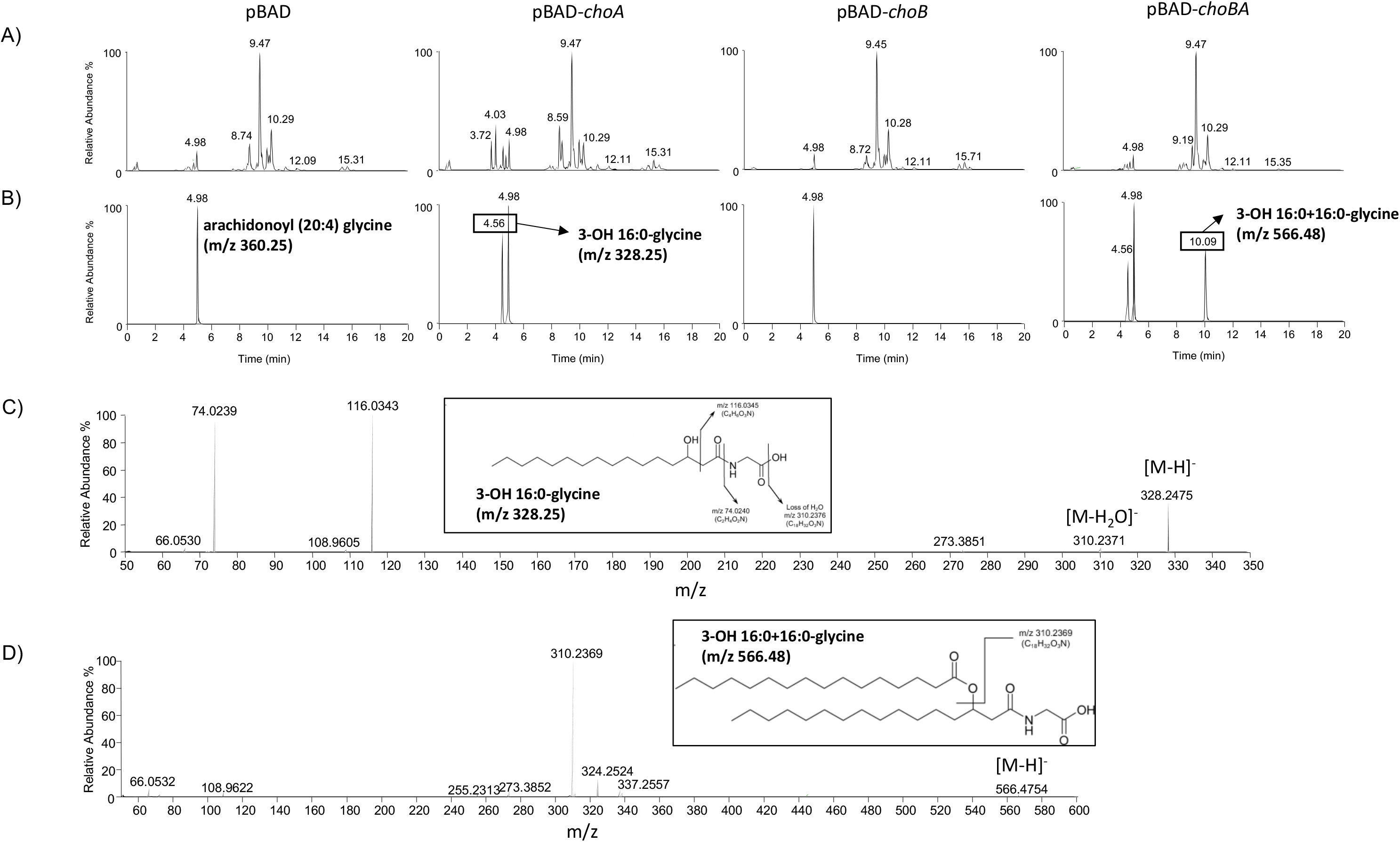
Identification of glycine lipids in E. coli overexpressing *choA* and *choB. E. coli* containing pBAD24, pBAD-*choA*, pBAD-*choB* or pBAD-*choBA* were cultured in the presence of 0.2% L-arabinose until mid-exponential phase and extracted lipids were analysed by LC-MS, as described in Materials and Methods. All samples were spiked with 500pmol arachidonyl (20:4) glycine as an internal standard (Rf=4.98 min, m/z 360.25). (A) The base peak intensity (BPI) chromatogram showing the range of lipids present in the *E. coli* membrane. All lipid profiles appear to be qualitatively similar, with the exception of pBAD-*choA* which shows increased peaks eluting with Rf of approx. 4 min where mono-acylated glycine molecules would be expected to be eluted. (B) Extracted ion chromatograms of peaks eluting with Rf values corresponding to mono- or di-acylated glycine species (for clarity only 3-OH 16:0 (Rf=4.56 min m/z 328.25) and 3-OH 16:0+16:0 (Rf=10.09 min m/z 566.48) are shown. The full list of identified molecules can be found in Table 1). (C) MS/MS fragmentation of the molecule eluting with Rf=4.56 confirming its identification as 3-OH 16:0-glycine. The same peak in both pBAD-*choA* and pBAD-*choBA* gave identical fragmentation profiles. (D) MS^3^ spectral analysis confirming the O-acylation of 3-OH 16:0 resulting in a structure of the m/z 566.48 compound that is consistent with a diacylated glycine (in this case 3-OH 16:0+16:0). All diacylated glycine molecules detected are listed in Table 1.

**Table 1.** Glycine lipids in *E. coli* expressing *choA* and *choB*

| Predicted size of acyl group (m/z) | Concentration (pmol/10<sup>9</sup> cells)±S.D. |  |  |
| --- | --- | --- | --- |
|  | vector | pBAD-*choA* | pBAD-*choAB* |
| 3-OH-12:0 | nd | 35.3±9.0 | nd |
| 3-OH-14:0 (300.2)* | nd | 4070.3±220.9 | 221.3±145.2 |
| 3-OH-14:1 (298.2)* | nd | 118.0±13.3 | nd |
| 3-OH-15:0 (314.2)* | nd | 52.4±14.4 | 9.2<sup>#</sup> |
| 3-OH-16:0 (328.2)* | nd | 3954.6±215.3 | 1584.9±843.4 |
| 3-OH-16:1 (326.2)* | nd | 6831.9±264.0 | 382.3±267.1 |
| 3-OH-18:0 (356.1)* | nd | 7.8±1.1 | 37.4±33.7 |
| 3-OH-18:1 (354.2)* | nd | 2539.2±227.5 | 1697.4±994.2 |
| Total monoacylated glycine |  | 17609.5±965.5 | 3923.3±2283.6 |
| 3-OH-14:0 + 12:0 (482.4)* | nd | nd | 436.6±228.5 |
| 3-OH-16:0+12:0/3-OH-14:0+14:0 (510.4)* | nd | nd | 5864.1±2999.9 |
| 3-OH-16:1+12:0 (508.4)* | nd | nd | 652.0±356.9 |
| 3-OH-29:0 | nd | nd | 240.5±122.1 |
| 3-OH-16:0+14:0 (538.4)* | nd | 16.3±5.7 | 17407.0±6125.0 |
| 3-OH-16:1+14:0/3-OH-16:0+14:1 (536.4)* | nd | nd | 5154.6±2257.9 |
| 3-OH-30:2 | nd | nd | 452.0±238.0 |
| 3-OH-16:0+15:0 (552.4)* | nd | nd | 458.8±228.3 |
| 3-OH-31:1 | nd | nd | 152.3±47.2<sup>#</sup> |
| 3-OH-16:0+16:0 (566.4)* | nd | 44.5±10.8 | 3901.3±141.9 |
| 3-OH-16:0+16:1/3-OH-18:1+14:0 (564.4)* | nd | 39.4±14.6 | 10673.8±3157.3 |
| 3-OH-16:1+16:1 (562.4)* | nd | nd | 1322.0±429.6 |
| 3-OH-33:1 | nd | nd | 131.4±74.3 |
| 3-OH-16:0+18:1/3-OH-18:1+16:0 (592.5)* | nd | 27.7±15.2 | 1540.3±23.7 |
| 3-OH-18:1+16:1 (590.4)* | nd | nd | 1275.5±218.8 |
| 3-OH-36:2 | nd | nd | 179.5±43.9 |
| Total diacylated glycine |  | 127.9±46.3 | 49841.7±16693.3 |
| Total acylated pool |  | 17737.4 | 53765.0 |

<sup>#</sup>: not detected in all biological replicates
\*: acylated glycine confirmed by MS/MS fragmentation. Acyl group designation is indicative, based on the predicted number of carbons in the lipid species.

### *GlsB is required for the production of commendamide in* B. thetaiotaomicron

We wanted to confirm that *glsA* and *glsB* were involved in the production of GL in *Bacteroides*. To do this we constructed a Δ*glsB* deletion mutation in *Bacteroides thetaiotaomicron* VPI-5482. We also constructed a strain whereby a native copy of *glsB* was inserted into the genome of the Δ*glsB* mutant strain (Δ*glsB::glsB*). Unfortunately, despite several attempts, we were unable to construct a knock-out mutation of *glsA* (or a double knock-out of *glsA glsB*) in *B. thetaiotaomicron*. Nonetheless, we were able to identify lipid species in both WT and the complemented Δ*glsB::glsB* strain that were consistent with commendamide and other N-acylated derivatives of glycine (see Figure 3 and Table 2). Further mass spectrometric analysis indicated the presence of diacylated GL in both WT and the complemented Δ*glsB::glsB* strain but not in the Δ*glsB* mutant (see Figure 3 and Table 3). These species showed differences in their retention time compared to the GL found in *E. coli* and gave rise to multiple, partially resolved chromatographic peaks. *Bacteroides* are known to produce branched-chain fatty acids although from our MS/MS analysis it was not possible to definitively assign whether the glycines were acylated with straight or branched-chain acyl groups (e.g. 16:0 or methyl-15:0) or if they were iso- or anteiso-branched (28). Importantly, we could not detect any acylated glycine in the Δ*glsB* mutant (see Table 2). Therefore, *glsB* is required for the production of all acylated glycine species in *B. thetaiotaomicron*. Interestingly, a comparison of the total lipid chromatogram did reveal both qualitative and quantitative differences between the Δ*glsB* mutant and both the WT and Δ*glsB::glsB* strain (see Figure 3A). Although a comprehensive analysis of these differences is not the objective of this study, we did determine that Lipid 654 is produced by *B. thetaiotaomicron* but absent from the Δ*glsB* mutant (see Supplementary Figure S1). In addition, a series of molecules with retention times approximately 14-16 mins and m/z ratios ranging from 1200-1300 were also completely absent from the Δ*glsB* mutant. The identity of these species is currently under investigation. Therefore, our data confirms that *glsB* is required for the production of GL and Lipid 654 in *B. thetaiotaomicron* and a mutation in *glsB* results in significant qualitative and quantitative changes in the lipid profile of the membranes of this bacterium.

**Figure 3.**
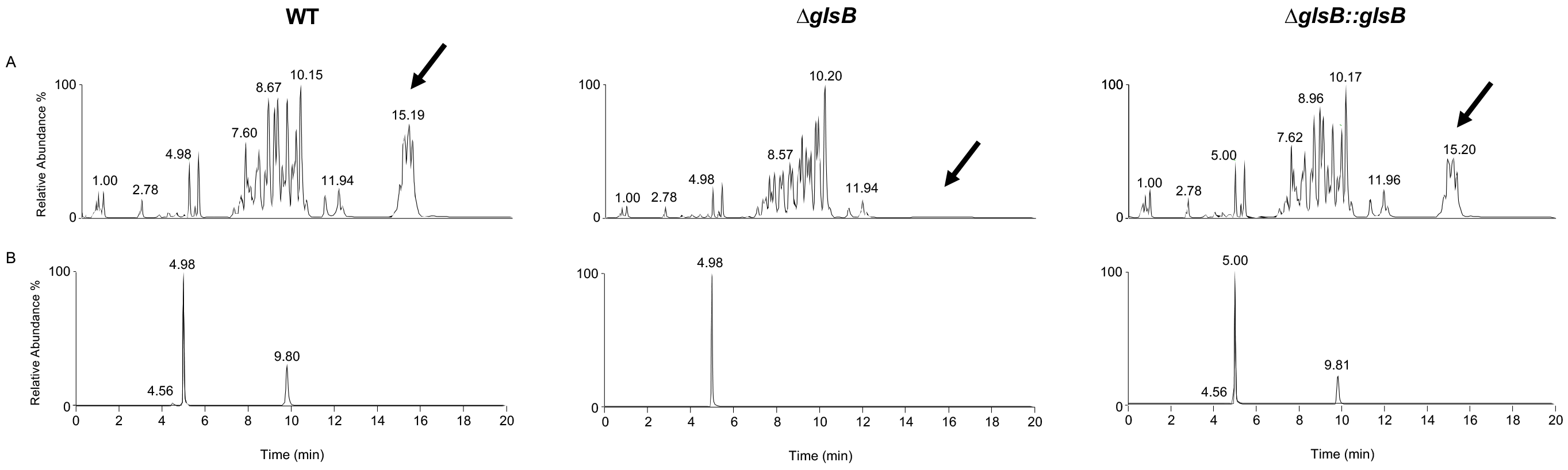
Identification of glycine lipids in *B. thetaiotaomicron*. Cells (as indicated) were cultured in BHIS broth until mid-exponential phase and extracted lipids were analysed by LC-MS, as described in Materials and Methods. All samples were spiked with 500pmol arachidonyl (20:4) glycine as an internal standard (Rf=4.98 min, m/z 360.25). (A) The base peak intensity (BPI) chromatogram showing the range of lipids present in the membrane of *B. thetaiotaomicron*. Clear qualitative (indicated by arrow) and quantitative differences are observed between the profiles of WT, ∆*glsB* and ∆*glsB::glsB* strains. (B) Extracted ion chromatograms of peaks eluting with Rf values corresponding to mono- or di-acylated glycine species. For clarity only 2 ions are shown corresponding to 3-OH 16:0 (Rf=4.56 min m/z 328.25) and C32:0 (Rf=9.8 min m/z 566.4). The full list of identified molecules can be found in Table 2.

**Table 2.** Glycine lipids in *B. thetaiotaomicron*

| Predicted size of acyl group (m/z) | Concentration (pmol/10<sup>9</sup> cells) ±S.D. |  |  |
| --- | --- | --- | --- |
|  | WT | ΔglsB | ΔglsB::glsB |
| 3-OH-15:0/3-OH-methyl 14:0 (314.2)* | 67.9±23.8<sup>#</sup> | nd | 64.6±16.9<sup>#</sup> |
| 3-OH-16:0/3-OH-methyl 15:0 (328.2)* | 252.1±36.3<sup>#</sup> | nd | 137.2±98.1 |
| 3-OH-17:0/3-OH-methyl 16:0 (342.2)* | 429.9±71.6<sup>#</sup> | nd | 223.0±157.0 |
| Total monoacylated glycine | 749.9±131.7 | 0 | 424.8±272 |
| 3-OH-methyl 15:0+13:0/3-OH-methyl 14:0+14:0 (524.4)* | 223.2±25.5 | nd | 169.4±30.9 |
| 3-OH-methyl 15:0+14:0/3-OH-methyl 14:0+15:0 (538.4)* | 792.1±81.2 | nd | 635.4±92.0 |
| 3-OH-methyl 15:0+15:0/3-OH-methyl 16:0+14:0 (552.4)* | 2000.3±139.3 | nd | 1372.0±181.3 |
| 3-OH-methyl 16:0+15:0/3-OH-methyl 15:0+16:0 (566.4)* | 2602.6±118.9 | nd | 1768.0±259.6 |
| Total diacylated glycine | 5618.2±364.9 | 0 | 3944.8±563.8 |
| **Total acylated pool** | **6368.1** | **0** | **4369.6** |

nd: not detected
<sup>#</sup>: only detected in 2 (out of 3) biological replicates
\*: acylated glycine confirmed by MS/MS fragmentation. Acyl group designation is indicative, based on the predicted number of carbons in the lipid species.

### *The expression of* glsB *and* glsA *is constitutive in* B. thetaiotaomicron

We wanted to examine the expression levels of *glsA* and *glsB* in *B. thetaiotaomicron* during growth. Cells were cultured to mid-exponential phase in either rich (BHIS) or defined (DMM) growth media and total RNA was isolated. RT-PCR analysis suggested that *glsA* and *glsB* are on different transcripts in *B. thetaiotaomicron* (see Figure 4A). To determine the expression profile of *glsA* and *glsB* we examined the data from previously published microarray experiments undertaken using *B. thetatiotaomicron* cultured under different *in vitro* and *in vivo* conditions (29, 30). In TYG broth, a rich growth medium composed of tryptone, yeast extract and glucose, the levels of expression of both *glsA* and *glsB* is highest during early exponential phase and the expression of both genes decreases over time (see Figure 4B). A similar trend is observed during growth in minimal media but the decrease in *glsA* and *glsB* expression over time is not as strong (see Figure 4B). Therefore, the expression of *glsA* and *glsB* may be linked to growth rate. Moreover, *glsA* and *glsB* are also expressed during colonization of the cecum of mice by *B. thetaiotaomicron* (see Figure 4B). Therefore, *glsA* and *glsB* appear to be constitutively expressed in *B. thetaiotaomicron* during growth *in vitro* and *in vivo*.

**Figure 4.**
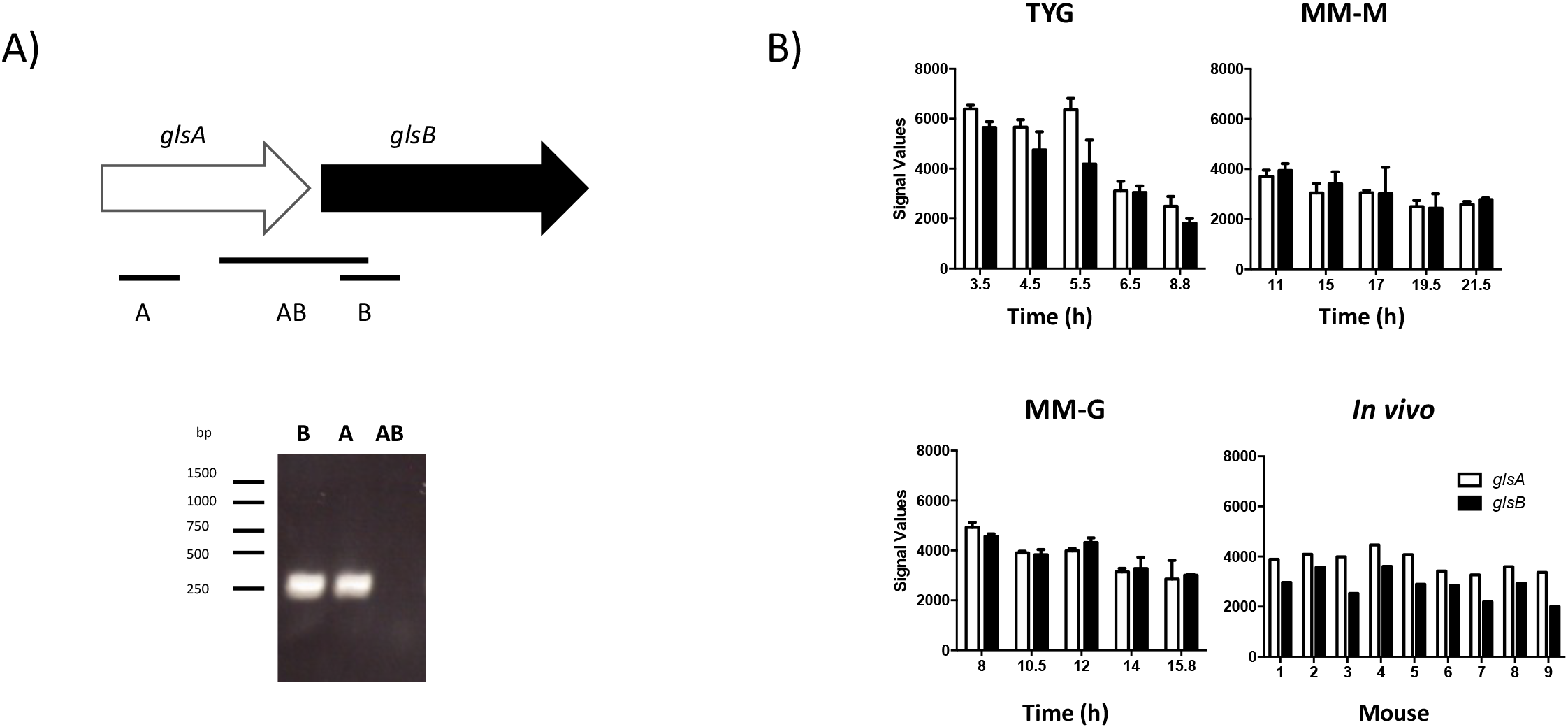
Analysis of the expression of *glsA* and *glsB* in *B. thetaiotaomicron*. (A) RT-PCR transcript analysis of expression from the *glsAB* locus. B. thetaiotaomicron cells were grown to mid-exponential phase in BHIS broth and RNA was extracted and back-transcribed into cDNA (as described in Materials and Methods). Transcript analysis was undertaken using primers combinations that amplify a region specific to *glsA* (A), *glsB* (B) or the intergenic region (AB). (B) Expression of *glsA* and *glsB* under different *in vitro* and *in vivo* growth conditions. Normalized microarray data was extracted from datasets available in the GEO database (Accession number: GSE2231). Sample preparation and analysis is described in (29, 30). For the *in vitro* samples, *B. thetaiotaomicron* was cultured in chemostats using different media (TYG broth, Minimal Medium-Maltose (MM-M), Minimal Medium-Glucose (MM-G)). At the indicated times cells were harvested, RNA was extracted and expression profiling was undertaken using custom *B. thetaiotaomicron* GeneChips. The data presented are the mean of 2 biological replicates and the error bars represent the standard deviation. For the *in vivo* experiments, individual germ-free NRBI mice (n=9) were monoassociated with *B. thetaiotaomicron* and fed a standard chow diet for 10 days. RNA was extracted from the cecal contents of each mouse and used for expression profiling using the custom *B. thetaiotaomicron* GeneChips.

### *The* glsB *gene is required for normal growth* in vitro

During preliminary experiments we observed that, when colonies were inoculated from agar plates into BHIS broth and incubated overnight, there was significantly reduced growth of the Δ*glsB* mutant compared to WT cultures. This suggested that *glsB* might be important for the normal growth of *B. thetaiotaomicron*. In order to quantify this observation we set up 10 overnights, from fresh BHIS agar plates inoculated with WT, the Δ*glsB* mutant or the Δ*glsB::glsB* strain and the cultures were incubated at 37°C for 18h at which point the final OD_600_ was taken as a measurement of growth. WT and Δ*glsB::glsB* cultures grown under these conditions reached OD_600_ values with a mean of 1.42 +/-0.042 and 1.25 +/-0.25, respectively (see Figure 5). However, the OD_600_ values of cultures inoculated with the Δ*glsB* mutant were significantly lower (0.36 +/-0.23; P<0.001) confirming that the Δ*glsB* mutant has a strong growth defect under these conditions. Interestingly, when Δ*glsB* mutant cells from the overnight broth cultures were inoculated into fresh broth cultures there was no observed defect in growth rate between the WT and the Δ*glsB* mutant (see Figure 5B). Therefore, our data suggests that the Δ*glsB* mutant is not defective in growth *per se* but *glsB* may be required to facilitate adaptation of *B. thetaiotaomicron* to the transition from growth on a solid surface to growth in liquid broth.

**Figure 5.**
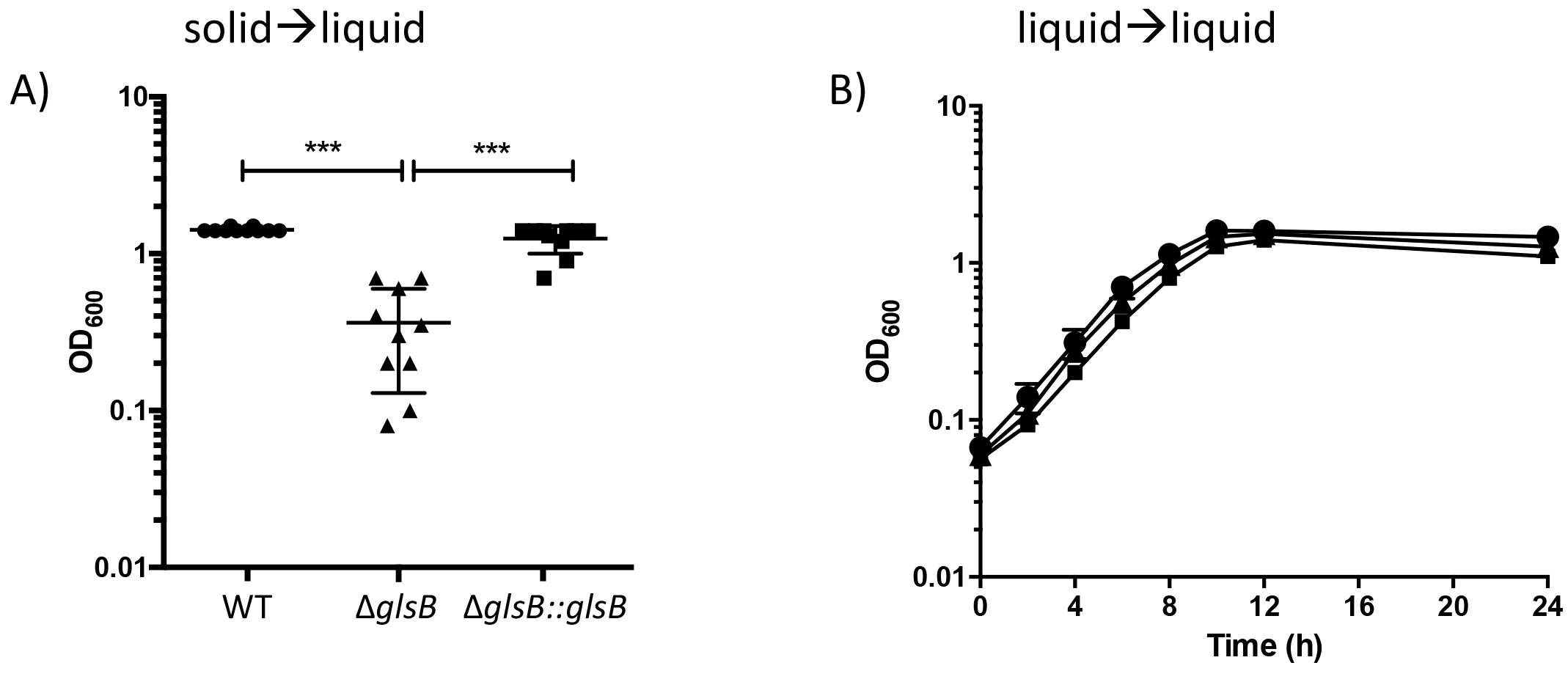
The *glsB* mutant is unable to normally transition from solid to liquid growth media. (A) *B. thetaiotaomicron* (WT, ∆*glsB* and ∆*glsB::glsB*) was grown on BHIS agar and individual colonies (n=10) were inoculated into fresh BHIS broth (solid→liquid). The cultures were incubated at 37°C for 24h and OD_600_ was used to measure growth. Each point represents an individual culture and the mean (+/-standard deviation) is presented (***P<0.0001 as determined using one-way ANOVA with Tukeys post-test for multiple comparisons). (B) Cells cultured in (A) were used to inoculate fresh BHIS broth and cells were grown at 37°C and OD_600_ was measured at the indicated time. Each strain was grown in triplicate and each point is the mean of the replicates and errors bars represent the standard deviation.

### *The* glsB *gene is required for adaptation to stress* in vitro

The transition from growth in solid to liquid media may represent a stress to the bacterium. Therefore, we decided to assess the sensitivity of the Δ*glsB* mutant to different stresses that would normally be encountered by *B. thetaiotaomicron* i.e. bile stress and the presence of oxygen in air. *B. thetaiotaomicron* was cultured in BHIS broth to mid-exponential phase before the cells were transferred to fresh medium supplemented with 1% (w/v) porcine bile and the cells were incubated, anaerobically, for a further 14 h. Under these conditions both WT and Δ*glsB::glsB* cultures reached a final cell density of 2.6 x 10^9^ cfu ml^−1^ and 2.7 x 10^9^ cfu ml^−1^ respectively (see Figure 6A). This is only marginally lower than the cell density achieved when cells are grown under the same conditions but in the absence of bile (2.98 x 10^9^ cfu ml^−1^ and 2.9 x 10^9^ cfu ml^−1^, respectively) and this reflects the high level of bile tolerance associated with the *Bacteroides* [Wexler, 2007]. In contrast the Δ*glsB* mutant only achieved a cell density of 2.7 x 10^5^ cfu ml^−1^ when cultured in the presence of 1% (w/v) porcine bile (in contrast to 2.7 x 10^9^ cfu ml^−1^ when cultured in the absence of bile). Therefore, the Δ*glsB* mutant is approximately 10^4^-fold more sensitive to porcine bile than the WT (see Figure 6A). Similarly, the Δ*glsB* mutant exhibited a 10-fold increased sensitivity to exposure to air for 14h compared to both the WT and Δ*glsB::glsB* strain (see Figure 6B). Therefore, the *glsB* gene is important in *B. thetaiotaoimicron* to allow adaptation to a variety of stresses including exposure to bile and air.

**Figure 6.**
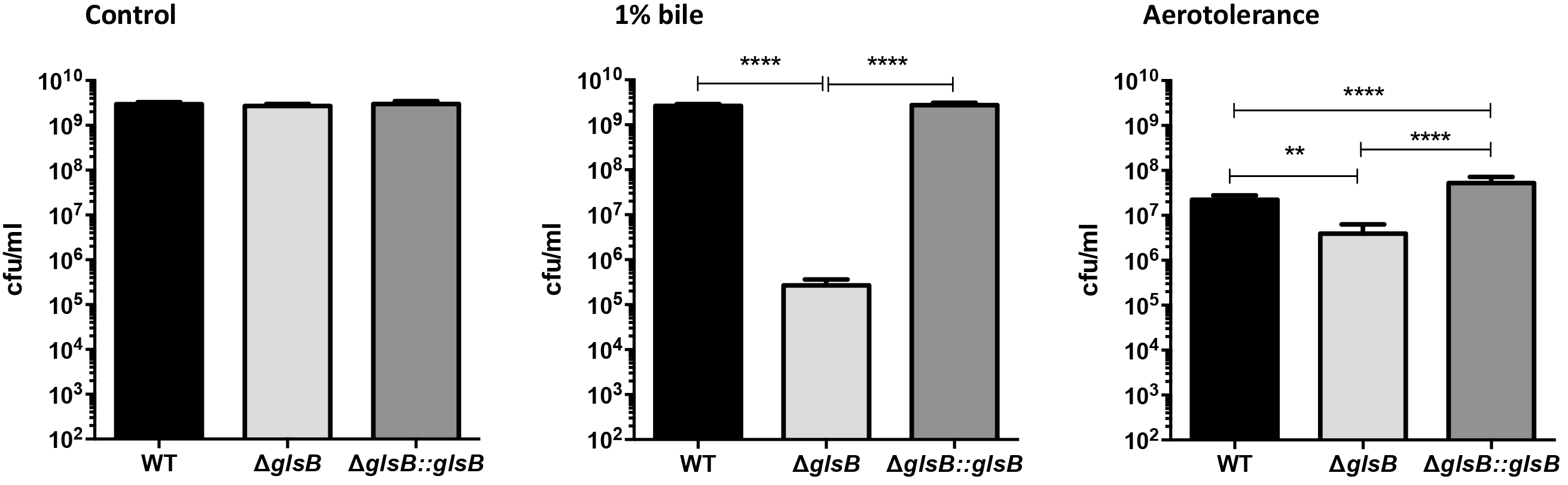
The *glsB* gene is required for adaptation to stress. *B. thetatiotaomicron* (WT, ∆*glsB* and ∆*glsB::glsB*) was cultured to mid-exponential phase (OD_600_=0.2) before (i) inoculation into BHIS broth followed by anaerobic incubation for 14h (Control); (ii) inoculation into BHIS broth supplemented with 1% (w/v) porcine bile followed by anaerobic incubation for 14h (1% bile); (iii) inoculation into BHIS broth followed by aerobic incubation (with vigorous shaking) for 14h (Aerotolerance). Viable cell counts were determined using serial dilutions that were plated onto BHIS agar followed by anaerobic incubation for 24-48h. The experiment was repeated 3 times and the error bars represent the standard deviation. Significance was determined using a one-way ANOVA with Tukeys post-test for multiple comparisons (**P<0.01, ***P<0.001, ****P<0.0001).

### *The* glsB *gene is required for normal colonization of the murine gut*

We wanted to determine the role, if any, of *glsB* during colonization of the mammalian gut. Therefore, germ-free C57BL/6 mice were subjected to a single, oral gavage of 10^8^ cfu of either WT *B. thetaiotaomicon* or Δ*glsB* mutant bacteria. Fecal pellets were collected on Day 2, 6, 9 and 12 post-gavage and bacteria were enumerated by viable plate counting on BHIS agar. On Day 2 the level of WT *B. thetaiotaomicron* in fecal pellets was 3.6 x 10^10^ cfu g-1 feces compared to a significantly lower level of 5.7 x 10^9^ cfu g^−1^ feces for the Δ*glsB* mutant (see Figure 7A). However, by Day 4, the Δ*glsB* mutant was found to be present in fecal pellets at the same level as the WT. Analysis of the cecal contents of mice collected on Day 14 indicated that there is a small, but significant, decrease in the level of the Δ*glsB* mutant in the cecum compared to WT *B. thetaiotaomicron* (see Figure 7B). Therefore, the Δ*glsB* mutant is affected in its ability to colonize the murine gut, particularly during the early stages of colonization.

**Figure 7.**
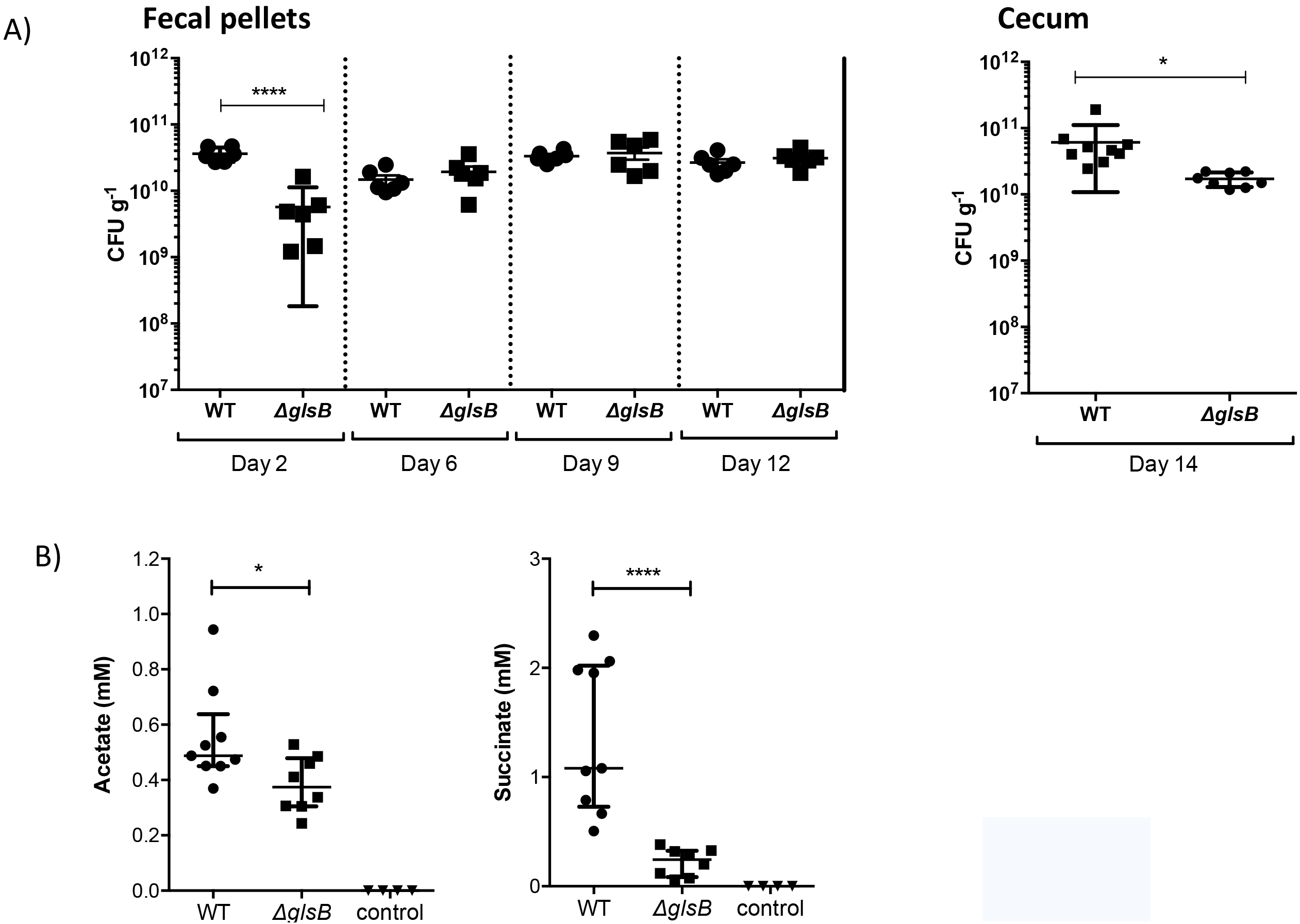
*B. thetatiotaomicron* requires the *glsB* gene for normal colonization of the gut of mice. Germ-free C57BL/6 mice were gavaged with a single 108 cfu dose of *B. thetaiotaomicron* WT (n=9) or ∆*glsB* mutant (n=8), as indicated. Control mice (n=4) were not colonized. (A) At the indicated time post-gavage fecal pellets were collected and bacteria were enumerated by plating fecal homogenates on BHIS agar. Bacteria were also enumerated in the cecum of colonized mice on Day 14 post-gavage. Control mice did not contain any bacteria at this point. Error bars represent the standard deviation of colonization levels in at least 5 mice (Unpaired t-test *P<0.05, ****P<0.001). (B) The contents of the cecum from each colonized and control mouse was examined for the presence of acetate and succinate by HPLC analysis. The error bars represent the 25-75% percentile values from the median and significance was determined using the Mann-Whitney test (*P<0.05, ****P<0.0001).

An important role for *B. thetaiotaomicron* in the gut is the conversion of dietary glycans into SCFA such as acetate and other organic acids e.g. succinate (4, 31). Therefore, we decided to use acetate and succinate production as a marker of *B. thetaiotaomicron* metabolism in the host. Using HPLC, we measured the level of acetate and succinate in the cecal contents collected from germ-free mice infected with either WT or Δ*glsB* mutant. As expected, we could not detect any acetate or succinate in the cecal contents of uninfected mice, confirming that these metabolites are exclusively derived from microbial activity in the gut (see Figure 7C). There was a small, but significant (Mann-Whitney test; *P*=0.036), decrease in the level of acetate present in the cecal contents of mice colonized with the Δ*glsB* mutant compared to WT (0.37mM (0.24-0.53mM) vs 0.49mM (0.37-0.94mM), respectively). The level of succinate was also significantly (Mann-Whitney test; *P*<0.0001) reduced in the cecal contents of mice infected with the Δ*glsB* mutant compared to the WT bacteria (0.25mM (0.06-0.38mM) vs 1.1mM (0.51-2.3mM), respectively). Therefore, the metabolism of the Δ*glsB* mutant is different from WT *B. thetaiotaoimicron* during growth in the murine gut.

## Discussion

The acylated amino acids, GL and flavolipin, have previously been identified in the membranes of several different members of the Phylum Bacteroidetes (8, 17, 32). In this study we have identified, for the first time, the genes required for the production of these acylated amino acids. Using genetics and high resolution LC/MS we show that *glsB* (BT_3459) encodes a glycine N-acyltrasferase that is required for the production of both GL and flavolipin in *B. thetaiotaomicron*. We also present evidence that *glsA* (BT_3458), a gene found immediately upstream from *glsB* on the *B. thetaiotaomicon* genome, encodes an O-acyltransferase that is required for the efficient production of GL. Interestingly, the overproduction of *glsA* and *glsB* in *E. coli* results in the synthesis of only GL suggesting the presence of another activity in *B. thetaiotaomicron* that may convert GL to flavolipin.

Flavolipin (i.e. Lipid 654) has been reported to signal to eukaryotic cells through an interaction with TLR2 (23, 33). The diacylated Lipid 654 has been shown to be converted into a more potent monoacylated derivative, Lipid 430 (or lyso-Lipid 654), through the action of phospholipase A2 activity in the host (17). N-acyl amino acids are important endogenous signaling molecules produced in the human host e.g. N-arachidonoyl glycine has been shown to block pain perception in mice (34-36). Genes encoding N-acyl synthases are enriched in the human gut microbiome and both Lipid 654 and Lipid 430 have been found in tissues that are distal from the gut indicating that these molecules can be distributed around the human host (17, 36). Therefore, there is accumulating evidence supporting an important role for the gut microbiota in the production of acylated amino acids, such as flavolipin, that can act as signaling molecules in the host.

During periods of phosphorous starvation some bacteria access cellular phosphorous reserves by using acylated amino acids such as OL to replace the phospholipids normally found in bacterial membranes (37-39). On the other hand, OL production in *Rhizobium tropici* and *Burholderia cepacia* has been shown to be important for the normal tolerance of the bacterial cell to acid and temperature stress (13, 40, 41). Our data suggests a role for GL/flavolipin during the response of *B. thetaiotaomicron* to different stresses. Therefore, a *glsB* deletion mutant was compromised in its ability to adapt to various stresses including transition from liquid to solid media, exposure to bile and exposure to air. Interestingly we could not construct a deletion in *glsA* suggesting that this gene might be essential in *B. thetaiotaomicron*. The reason for this essentiality is unclear but it might suggest that *glsA* is involved in the O-acylation of additional lipid species. The orthologues of *glsA* and *glsB* have been successfully deleted in *Bacteroides fragilis* suggesting that the essentiality of *glsA* might be strain-dependent (42). In *B. thetaiotaomicron* we have shown that there are both qualitative and quantitative changes in the lipid profile of the *glsB* mutant suggesting that the cells do compensate for the absence of GL and flavolipin. Therefore, it is possible that *glsA* is required during the compensatory changes that occur in the absence of GL/flavolipin in *B. thetaiotaomicron* although this remains to be determined.

In this study we show that the Δ*glsB* mutant in *B. thetaiotaomicron* is affected in its ability to colonize the gut of a GF mouse. A deletion of both the *glsA* and *glsB* orthologues in *B. fragilis* (named *hlyB* and *hlyA*, respectively) was also attenuated for virulence in a mouse abscess model supporting a role for these genes during in vivo growth (43). We have shown that, in contrast to the WT, the Δ*glsB* mutant produces decreased levels of both acetate and succinate whilst in the cecum indicating that there are differences in the metabolism between the WT and the Δ*glsB* mutant. Acetate and succinate are important end-products of carbohydrate metabolism in *B. thetaiotaomicron*. Acetate is produced from acetyl-CoA via the acetate kinase (AckA)-phosphate acetyltransferase (Pta) pathway resulting in the generation of ATP. The reduction in acetate production in the Δ*glsB* mutant is small and may not by physiologically important. Succinate production is via phosphoenolpyruvate, oxaloacetate, malate and fumarate and is linked to the production of reducing equivalents (through the regeneration of NAD(P)+) and the formation of a proton motive force. Therefore, the significantly reduced levels of succinate produced by the Δ*glsB* mutant would be expected to compromise metabolic flux and, therefore, reduce fitness in the gut. In support of this, Δ*glsB* (BT_3459) has been identified as an essential determinant for the colonization of the GF mouse gut during an INSeq screen with *B. thetaiotaomicron* (44).

Therefore, we have shown that *glsB*, and presumably the production of GL and/or flavolipin, is an important fitness factor in *Bacteroides*, required for the adaptation to stress and normal colonization of the mammalian gut, particularly in the presence of a competing microbiota.

## Acknowledgements

This work was funded by an Investigator Award from Science Foundation Ireland (SFI) to D.J.C. 912/IP/1493) and by funding received through APC Microbiome Ireland, a research institute supported by SFI (SFI/12/RC/2273). S.R.T., M.K.D. and P.D.W. gratefully acknowledge the financial support of the European Regional Development Fund, Scottish Funding Council and Highlands and Islands Enterprise.

## Author contributions

D.J.C. conceived the study and A.L. carried out all of the experiments except the lipidomics. P.D.W., S.R.T and M.K.D. carried out all lipidomic experiments and analyses. A.L. and D.J.C. analysed the data and D.J.C. wrote the manuscript with the help of A.L. and P.D.W. All authors reviewed the manuscript.

## Bibliography

1. Qin J, Li R, Raes J, Arumugam M, Burgdorf KS, Manichanh C, Nielsen T, Pons N, Levenez F, Yamada T, Mende DR, Li J, Xu J, Li S, Li D, Cao J, Wang B, Liang H, Zheng H, Xie Y, Tap J, Lepage P, Bertalan M, Batto J-M, Hansen T, Le Paslier D, Linneberg A, Nielsen HB, Pelletier E, Renault P, Sicheritz-Ponten T, Turner K, Zhu H, Yu C, Li S, Jian M, Zhou Y, Li Y, Zhang X, Li S, Qin N, Yang H, Wang J, Brunak S, Doré J, Guarner F, Kristiansen K, Pedersen O, Parkhill J, Weissenbach J, MetaHIT Consortium, Bork P, Ehrlich SD, Wang J. 2010. A human gut microbial gene catalogue established by metagenomic sequencing. Nature 464:59–65.

2. Grondin JM, Tamura K, Déjean G, Abbott DW, Brumer H. 2017. Polysaccharide Utilization Loci: Fueling Microbial Communities. J Bacteriol 199:e00860–16–15.

3. Porter NT, Martens EC. 2017. The Critical Roles of Polysaccharides in Gut Microbial Ecology and Physiology. Annu Rev Microbiol 71:349–369.

4. Wexler AG, Goodman AL. 2017. An insider’s perspective: *Bacteroides* as a window into the microbiome. Nature Microbiol 2:17026.

5. Marcobal A, Barboza M, Sonnenburg ED, Pudlo N, Martens EC, Desai P, Lebrilla CB, Weimer BC, Mills DA, German JB, Sonnenburg JL. 2011. *Bacteroides* in the Infant Gut Consume Milk Oligosaccharides via Mucus-Utilization Pathways. Cell Host Microbe 10:507–514.

6. Hill CJ, Lynch DB, Murphy K, Ulaszewska M, Jeffery IB, O’Shea CA, Watkins C, Dempsey E, Mattivi F, Tuohy K, Ross RP, Ryan CA, O’Toole PW, Stanton C. 2017. Evolution of gut microbiota composition from birth to 24 weeks in the INFANTMET Cohort. Microbiome 5:4.

7. Geiger O, González-Silva N, López-Lara IM, Sohlenkamp C. 2010. Amino acid-containing membrane lipids in bacteria. Prog Lipid Res 49:46–60.

8. Sohlenkamp C, Geiger O. 2016. Bacterial membrane lipids: diversity in structures and pathways. FEMS Microbiol Rev 40:133–159.

9. Vences-Guzmán MÁ, Geiger O, Sohlenkamp C. 2012. Ornithine lipids and their structural modifications: from A to E and beyond. FEMS Microbiol Lett 335:1–10.

10. Gao J-L, Weissenmayer B, Taylor AM, Thomas-Oates J, López-Lara IM, Geiger O. 2004. Identification of a gene required for the formation of lyso-ornithine lipid, an intermediate in the biosynthesis of ornithine-containing lipids. Mol Microbiol 53:1757–1770.

11. Weissenmayer B, Gao J-L, López-Lara IM, Geiger O. 2002. Identification of a gene required for the biosynthesis of ornithine-derived lipids. Mol Microbiol 45:721–733.

12. Vences Guzmán MÁ, Guan Z, Escobedo Hinojosa WI, Bermúdez Barrientos JR, Geiger O, Sohlenkamp C. 2015. Discovery of a bifunctional acyltransferase responsible for ornithine lipid synthesis in *Serratia proteamaculans*. Environ Microbiol 17:1487–1496.

13. Vences-Guzmán MÁ, Guan Z, Ormeño-Orrillo E, González-Silva N, López-Lara IM, Martínez-Romero E, Geiger O, Sohlenkamp C. 2011. Hydroxylated ornithine lipids increase stress tolerance in *Rhizobium tropici* CIAT899. Mol Microbiol 79:1496–1514.

14. Hölzl G, Sohlenkamp C, Vences Guzmán MÁ, Gisch N. 2018. Headgroup hydroxylation by OlsE occurs at the C4 position of ornithine lipid and is widespread in Proteobacteria and Bacteroidetes. Chem Phys Lipid 213:32–38.

15. Escobedo Hinojosa WI, Vences Guzmán MÁ, Schubotz F, Sandoval-Calderón M, Summons RE, López-Lara IM, Geiger O, Sohlenkamp C. 2015. OlsG (Sinac_1600) Is an Ornithine Lipid N-Methyltransferase from the Planctomycete *Singulisphaera acidiphila*. J Biol Chem 290:15102–15111.

16. Vences Guzmán MÁ, Guan Z, Bermúdez Barrientos JR, Geiger O, Sohlenkamp C. 2013. *Agrobacteria* lacking ornithine lipids induce more rapid tumour formation. Environ Microbiol 15:895–906.

17. Nemati R, Dietz C, Anstadt EJ, Cervantes J, Liu Y, Dewhirst FE, Clark RB, Finegold S, Gallagher JJ, Smith MB, Yao X, Nichols FC. 2017. Deposition and hydrolysis of serine dipeptide lipids of Bacteroidetes bacteria in human arteries: relationship to atherosclerosis. J Lipid Res 58:1999–2007.

18. Kawazoe R, Okuyama H, Reichardt W, Sasaki S. 1991. Phospholipids and a novel glycine-containing lipoamino acid in *Cytophaga johnsonae* Stanier strain C21. J Bacteriol 173:5470–5475.

19. Kawazoe R, Monde K, Reichardt W, Okuyama H. 1992. Lipoamino acids and sulfonoplipids in *Cytophaga johnsonae* Stanier strain C21 and six related species of *Cytophaga*. Arch Microbiol 158:171–175.

20. Kawai Y, Yano I, Kaneda K. 1988. Various kinds of lipoamino acids including a novel serine-containing lipid in an opportunistic pathogen *Flavobacterium*. Their structures and biological activities on erythrocytes. Eur J Biochem 171:73–80.

21. Farrokhi V, Nemati R, Nichols FC, Yao X, Anstadt E, Fujiwara M, Grady J, Wakefield D, Castro W, Donaldson J, Clark RB. 2013. Bacterial lipodipeptide, Lipid 654, is a microbiome-associated biomarker for multiple sclerosis. Clin Transl Immunology 2:e8.

22. Clark RB, Cervantes JL, Maciejewski MW, Farrokhi V, Nemati R, Yao X, Anstadt E, Fujiwara M, Wright KT, Riddle C, La Vake CJ, Salazar JC, Finegold S, Nichols FC. 2013. Serine Lipids of *Porphyromonas gingivalis* Are Human and Mouse Toll-Like Receptor 2 Ligands. Infect Immun 81:3479–3489.

23. Wang Y-H, Nemati R, Anstadt E, Liu Y, Son Y, Zhu Q, Yao X, Clark RB, Rowe DW, Nichols FC. 2015. Serine dipeptide lipids of *Porphyromonas gingivalis* inhibit osteoblast differentiation: Relationship to Toll-like receptor 2. Bone 81:654–661.

24. Lynch A, Crowley E, Casey E, Cano R, Shanahan R, McGlacken G, Marchesi JR, Clarke DJ. 2017. The *Bacteroidales* produce an N-acylated derivative of glycine with both cholesterol-solubilising and hemolytic activity. Sci Rep 7:1–10.

25. Cohen LJ, Kang H-S, Chu J, Huang Y-H, Gordon EA, Reddy BVB, Ternei MA, Craig JW, Brady SF. 2015. Functional metagenomic discovery of bacterial effectors in the human microbiome and isolation of commendamide, a GPCR G2A/132 agonist. Proc Natl Acad Sci (USA) 112:E4825–34.

26. Koropatkin NM, Martens EC, Gordon JI, Smith TJ. 2008. Starch catabolism by a prominent human gut symbiont is directed by the recognition of amylose helices. Structure 16:1105–1115.

27. Huda-Faujan N, Abdulamir AS, Fatimah AB, Anas OM, Shuhaimi M, Yazid AM, Loong YY. 2010. The impact of the level of the intestinal short chain Fatty acids in inflammatory bowel disease patients versus healthy subjects. Open Biochem J 4:53–58.

28. Mayberry WR. 1980. Hydroxy fatty acids in *Bacteroides* species: D-(--)-3-hydroxy-15-methylhexadecanoate and its homologs. J Bacteriol 143:582–587.

29. Sonnenburg JL, Xu J, Leip DD, Chen C-H, Westover BP, Weatherford J, Buhler JD, Gordon JI. 2005. Glycan foraging *in vivo* by an intestine-adapted bacterial symbiont. Science 307:1955–1959.

30. Sonnenburg ED, Sonnenburg JL, Manchester JK, Hansen EE, Chiang HC, Gordon JI. 2006. A hybrid two-component system protein of a prominent human gut symbiont couples glycan sensing in vivo to carbohydrate metabolism. Proc Natl Acad Sci USA 103:8834–8839.

31. Louis P, Flint HJ. 2017. Formation of propionate and butyrate by the human colonic microbiota. Environ Microbiol 19:29–41.

32. Moore EK, Hopmans EC, Rijpstra WIC, Villanueva L, Sinninghe Damsté JS. 2016. Elucidation and identification of amino acid containing membrane lipids using liquid chromatography/high-resolution mass spectrometry. Rapid Commun Mass Spectrom 30:739–750.

33. Olsen I, Nichols FC. 2018. Are Sphingolipids and Serine Dipeptide Lipids Underestimated Virulence Factors of *Porphyromonas gingivalis*? Infect Immun 86. doi: 10.1128/IAI.00035-18

34. Connor M, Vaughan CW, Vandenberg RJ. 2010. N-Acyl amino acids and N-acyl neurotransmitter conjugates: neuromodulators and probes for new drug targets. British J Pharmacol 160:1857–1871.

35. Bradshaw HB, Rimmerman N, Hu SSJ, Burstein S, Walker JM. 2009. Novel Endogenous N-Acyl Glycines: Identification and Characterization. Vitamins & Hormones 81:191–205.

36. Cohen LJ, Esterhazy D, Kim S-H, Lemetre C, Aguilar RR, Gordon EA, Pickard AJ, Cross JR, Emiliano AB, Han SM, Chu J, Vila-Farres X, Kaplitt J, Rogoz A, Calle PY, Hunter C, Bitok JK, Brady SF. 2017. Commensal bacteria make GPCR ligands that mimic human signalling molecules. Nature 549:48–53.

37. Barbosa LC, Goulart CL, Avellar MM, Bisch PM, Kruger von WMA. 2018. >Accumulation of ornithine lipids in *Vibrio cholerae* under phosphate deprivation is dependent on VC0489 (OlsF) and PhoBR system. Microbiology 164:395–399.

38. Diercks H, Semeniuk A, Gisch N, Moll H, Duda KA, Hölzl G. 2015. Accumulation of Novel Glycolipids and Ornithine Lipids in *Mesorhizobium loti* under Phosphate Deprivation. J Bacteriol 197:497–509.

39. Lewenza S, Falsafi R, Bains M, Rohs P, Stupak J, Sprott GD, Hancock REW. 2011. The *olsA* gene mediates the synthesis of an ornithine lipid in *Pseudomonas aeruginosa* during growth under phosphate-limiting conditions, but is not involved in antimicrobial peptide susceptibility. FEMS Microbiol Lett 320:95–102.

40. Taylor CJ, Anderson AJ, Wilkinson SG. 1998. Phenotypic variation of lipid composition in *Burkholderia cepacia*: a response to increased growth temperature is a greater content of 2-hydroxy acids in phosphatidylethanolamine and ornithine amide lipid. Microbiology 144:1737–1745.

41. González-Silva N, López-Lara IM, Reyes-Lamothe R, Taylor AM, Sumpton D, Thomas-Oates J, Geiger O. 2011. The dioxygenase-encoding *olsD* gene from *Burkholderia cenocepacia* causes the hydroxylation of the amide-linked fatty acyl moiety of ornithine-containing membrane lipids. Biochemistry 50:6396–6408.

42. Robertson KP, Smith CJ, Gough AM, Rocha ER. 2006. Characterization of *Bacteroides fragilis* hemolysins and regulation and synergistic interactions of HlyA and HlyB. Infect Immun 74:2304–2316.

43. Lobo LA, Jenkins AL, Jeffrey Smith C, Rocha ER. 2013. Expression of *Bacteroides fragilis* hemolysins *in vivo* and role of HlyBA in an intra-abdominal infection model. Microbiologyopen 2:326–337.

44. Goodman AL, McNulty NP, Zhao Y, Leip D, Mitra RD, Lozupone CA, Knight R, Gordon JI. 2009. Identifying genetic determinants needed to establish a human gut symbiont in its habitat. Cell Host Microbe 6:279–289.

